# Experimental Evolution of the Megaplasmid pMPPla107 in *Pseudomonas stutzeri* Enables Identification of Genes Contributing to Sensitivity to an Inhibitory Agent

**DOI:** 10.1101/538629

**Authors:** Brian A. Smith, Kevin Dougherty, Meara Clark, David A. Baltrus

## Abstract

Horizontally transferred elements such as plasmids can, at times, burden host cells with various metabolic and fitness costs. Our previous work demonstrated that acquisition of the *Pseudomonas syringae* megaplasmid pMPPla107 causes sensitivity to a growth inhibiting substance that is produced in cultures during growth under standard laboratory conditions. After 500 generations of laboratory passage of *P. stutzeri* lines containing pMPPla107, two out of six independent lines displayed resistance to this inhibitory agent. We therefore sequenced the genomes of isolates from each independent evolutionary line to identify the genetic basis of this resistance phenotype through comparative genomics. Our analysis demonstrates that two different compensatory mutations on the megaplasmid ameliorate the sensitivity phenotype: 1) a large deletion of approximately 368kb in pMPPla107 and 2) a SNP in the gene we name *skaA* for Supernatant Killing Activity. These results provide further evidence that costs associated with horizontal gene transfer can be compensated through single mutational events and emphasize the power of experimental evolution and resequencing to better understand the genetic basis of evolved phenotypes.

## INTRODUCTION

Plasmids are secondary replicons that can rapidly move across bacterial genomes increasing genomic plasticity through a process known as horizontal gene transfer (HGT). Thousands of genes can be transferred via HGT in an instance allowing for the colonization of new niches by the acquisition of genes encoding for metabolism, antibiotic resistance, virulence factors, and symbiosis thus enabling colonization of new niches(1–6). While plasmids could provide advantages for a bacterial cell in a given environment, horizontal gene transfer also brings many costs that may be manifested in phenotypic changes rather than lowered fitness alone(7–11). Outside of a handful of examples, relatively little is known about general trends underlying the mechanistic basis of such costs(8, 9, 12, 13).

We have previously shown that acquisition of the *Pseudomonas syringae* megaplasmid pMPPla107 by *Pseudomonas stutzeri* sensitizes this strain background to the presence of an inhibitory agent that has bacteriostatic properties (9, 10, 14). Sensitivity is found in pMPPla107’s native strain *P. syringae pv.* lachrymans 107 and can be transferred to various *Pseudomonas spp.* upon their acquisition of pMPPla107; thus, indicating the phenotype is linked to pMPPla107. Furthermore, production of the inhibitory agent is conserved across *Pseudomonas spp.* appears linked to Pseudomonas metabolism, and may be associated with an essential gene(14).

Although acquisition of plasmids by new host backgrounds often creates metabolic, physiological, and fitness costs, previous research has shown that various types of compensatory mutations occur rapidly on either the chromosome or plasmid and that such amelioration of costs enables the persistence of plasmids(15–19). Evolutionary experiments of *P. fluorescens* with the mercury resistant pQBR103 start with fitness costs in *P. fluorescens*. However, after hundreds of generations compensatory mutations in *gacA/gacS* occur in strains with and without selection using mercury, suggesting that a plasmid can exhibit parasitic behaviors influencing chromosomal mutations without any selective benefit to establish stable cohabitation(15). Furthermore, mutations in two helicases and an RNA polymerase subunit resulted in host dependence on the plasmid RP4 while also increasing the uptake of additional plasmids(16). Therefore, compensatory mutations may not only explain plasmid persistence mechanisms, but also increased plasmid promiscuity. For these reasons and to better understand the genetic basis of previously described phenotypic costs associated with pMPPla107(9), we carried out experimental evolution of *P. stutzeri* under conditions that selected for maintenance of pMPPla107. Our goal with these passage experiments was to identify strain backgrounds that have ameliorated known costs of pMPPla107 carriage with the hope that identification of compensatory mutations would provide better understanding of the genetic basis of these costs(14)

Here we resequence genomes from single colony isolates sampled from six independently evolving lines of *P. stutzeri* containing pMPPla107 after 500 generations of passage. Two of these lines evolved resistance to a well characterized, but currently unknown, inhibitory agent produced by Pseudomonas species. We further find that two different compensatory mutations provide resistance to this inhibitory agent and that both changes were found on the megaplasmid itself. Although one of these mutations was a large deletion that eliminated many different genes, the other was a single non-synonymous nucleotide polymorphism (SNP) that occurred in a gene with no known function Our work provides insights into the genetic basis of mutations that ameliorate fitness costs associated with plasmid acquisition but also more specifically inform our understanding of the genetic basis of sensitization of Pseudomonas strains to a currently unidentified inhibitory agent associated with maintenance of pMPPla107.

## METHODS

### Long Term Evolution Experiment

Six single colonies of *P. stutzeri* strain DBL408 were picked after growth on Salt Water LB (SWLB) agar(20), into independent 2mL cultures of SWLB liquid containing rifampicin (50ng/μL) and tetracycline (10ng/μL) within 5mL polypropylene tubes with caps. These cultures were grown within shaking (220rpm) at 27°C for two days at which point a subset of this culture was frozen in 40% glycerol at −80°C and labeled as “passage 0” while a 1:1000 (cells:media) dilution was also made into fresh 2mL of SW-LB. Each passage, cells were plated to SW-LB agar plates containing rifampicin (50ng/μL) and tetracycline (10ng/μL) to observe colony morphology in case of contamination. Tetracycline in the media selected for maintenance of the megaplasmid in strain DBL408. Every 10 passages, a 750μL sample of the culture was mixed with 40% final concentration of glycerol and stored at −80°C. This process was repeated for approximately 500 generations of growth (Log2 of 1000=9.96 divisions per passage; 50 passages total).

### Genome Sequencing and Annotation

Single colonies of each generation 500 line were picked and grown overnight in 2mL of SW-LB with rifampicin (50ng/μL) and tetracycline (10ng/μL). DNA samples were extracted from these cultures using a Promega Wizard kit. *P. stutzeri* lines 1B, 4B, 5B, and 6B were sequenced using 100bp paired end reads on an Illumina HiSeq (SRA in progress). *P. stutzeri* lines 2B and 3B were sequenced using 250bp paired end reads on an Illumina MiSeq by MicrobesNg (SRA in progress). We used Prokka(21) gene annotations of pMPPla107 from a previous publication(5) and the annotations from the *P. stutzeri* 28a24 reference sequence (Accession: CP007441.1).

### Mapping Reads and Calling Variants

Illumina reads from all six evolved lines were mapped to the *P. stutzeri* 23a24 and pMPPla107 references (Accession No.: CP007441 and NZ_CP031226.1 respectively) using the Geneious11.1.3 (https://www.geneious.com/) mapper. Parameters used for the mapping step were: do not trim, gaps allowed, maximum gap per read = 10%, word length = 18, ignore words repeated more than 12 times, maximum mismatches per read = 20%, maximum gap size = 15, index word length = 13, maximum ambiguity = 4, accurately map reads with errors to repeat regions.

Additionally the sensitivity parameter was set at medium-low sensitivity with up to 5 iterations. We found that mapping at higher sensitivities did not change our outputs and therefore chose this setting. After mapping, variants were called inside and outside of coding regions with a conservative frequency filter of 0.90. Variant maximum P-values were set at 10^−6^ and a minimum strand-bias P-value of 10^−5^ was also used. Since we were interested in gene mutations only in 5B responsible for resistance to the inhibitory agent we filtered for unique SNPs by removing redundant SNPs that occurred in > 1 evolved lines. This gave us a set of candidate genes to then conduct genetic analyses, allowing us to confirm a causative gene for the sensitivity phenotype. In cases where genes appeared to have higher rates of variance, we pruned our SNP data by removing variant calls in high variance regions with less than 30x coverage.

### Synteny Plots

SynMap is a web-based software found at genomeevolution.org used to build synteny plots of sequence data(22). We used SynMap2 with the LAST algorithm and default parameters to compare the sequences of ancestral pMPPla107 and pMPPla107-4B(22). DAGChainer Options were: nucleotide distance, −D = 20, and −A = 5. Tandem duplication distance was set to 10 and the C-score was set to 0.

### Inhibitory Agent Sensitivity Test

We followed previous protocols to test the inhibitory agent against the six evolved lines, as described elsewhere(9, 14). Briefly, overlays were prepared by mixing cells grown for four hours with 0.4% molten agar and plated on to KB plates. Overlays were allowed to solidify for approximately 15min. The inhibitory agent was collected by growing *P. stutzeri* for 24-48hrs, centrifuging cells at 10,000 × *g* for 5min, and sterilizing supernatants through a 0.22μm filter. After sterilization 10μL of supernatants were spotted onto the overlay plate and allowed to dry. Overlay plates were grown at 27°C for approximately 24hrs at which point, zones of inhibition were observed.

### Conjugation of Evolved Megaplasmids into Ancestral P. stutzeri

Ancestral *P. stutzeri* (DBL386) was used with either evolved lines 4B or 5B to conduct a biparental conjugation by mixing 1:1 mixture of overnight cultures. Mixed cells were centrifuged at 3000 × *g* for 3min and supernatants were removed without disturbing the pellet. Pellets were washed and resuspended in 1mL of 10mM MgCl_2_. Centrifugation and washing steps were repeated once more. 10μL and 100μL of resuspended cells were spread and for 24-48hr at 27°C on KB plates with rifampicin (50ng/μL) and tetracycline (10ng/μL). Resistant colonies underwent diagnostic PCR for presence of pMPPla107 using primers from Baltrus et al. 2011(23).

### Gene function prediction with Phyre2 and blastx

We attempted to predict functional characteristics of *skaA* using the Phyre2 web server. The amino acid for *skaA* was used as input and the intensive setting was selected(24). We also used the nucleotide sequence of *skaA* as input into the NCBI non-redundant protein sequences BLAST database using blastx using the BLOSUM62 matrix, expect threshold = 10, word size = 6, and max target sequences = 100. (date of search last search: January 21^st^ 2019)(25).

## RESULTS

### Genome Sequencing Reveals 2 of 6 Evolved Lines Gain Resistance to a Previously Described Inhibitory Agent

Given our interest in a phenotype involving sensitization of Pseudomonas strains to an unknown inhibitory agent after acquisition of pMPPla107(9), we screened for the presence of inhibition in these evolved lines. Single colony isolates from two out of six lines (referred to from here on as DBL408-4BGen500 and DBL408-5BGen500) revert to the non-pMPPla107 phenotype and demonstrate resistance to this inhibitory agent (Figure 1).

**Figure 1:**
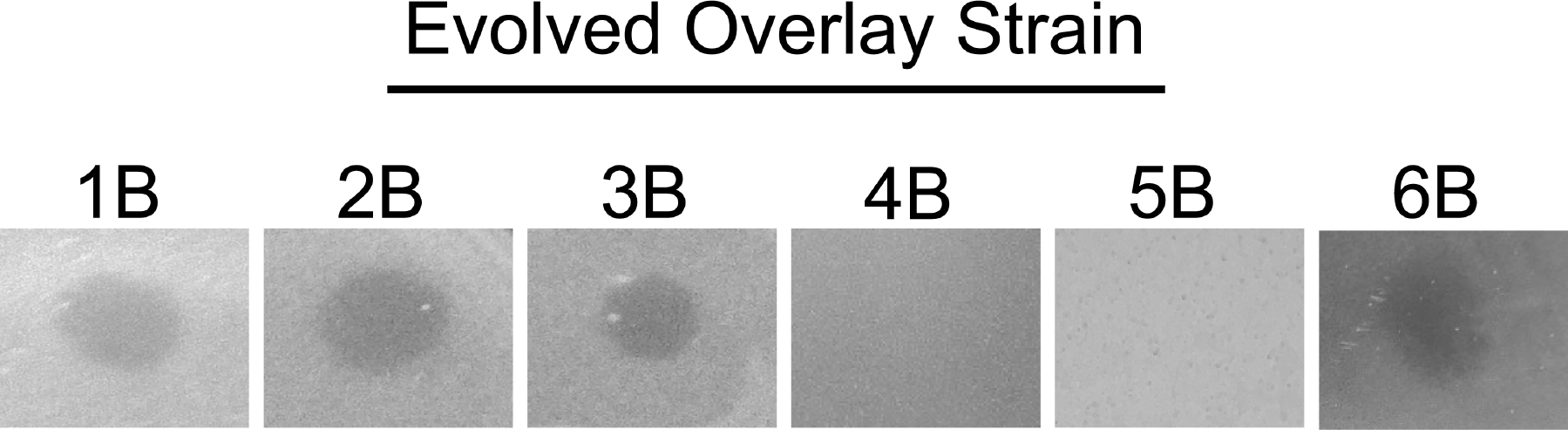
Testing inhibitory agent on all six evolved lines reveals two resistant lines. Six lines carrying pMPPla107 were evolved for 500 generations and tested for sensitivity against inhibitory agent found in *Pseudomonas spp.* supernatants. Lines 4B and 5B revert to a non-pMPPla107 phenotype where a zone of inhibition is not present indicating resistance to the inhibitory agent. All overlays were plated after 4 hours of growth n KB and spotted with 10μL of *P. stutzeri* filter sterilized supernatants. The number (1, 2, 3…) indicates the individual lines and B indicates the second of two isolates taken at Generation 500. All images are representative of three biological replicates.

In previous evolutionary studies focusing on plasmids, the burden of plasmid acquisition resulted in compensatory mutations present on host chromosomes(15, 16, 26, 27). To identify where the resistance mutations occurred, we analyzed the genomes of six laboratory passage strains of *P. stutzeri* after 500 generations under conditions that selected for maintenance of megaplasmid pMPPla107. Sequencing of the chromosome and pMPPla107 revealed variants across all six evolved lines (Tables 1 and 2 DOIs: doi.org/10.6084/m9.figshare.7393415 and doi.org/10.6084/m9.figshare.7393493 respectively), the majority of which (209/219) occur on the chromosome. All six evolved *P. stutzeri* lines also have a SNP on pMPPla107 at 731,508bp in *qseF* indicating this was a mutation either occurred prior to the start of the evolutionary experiment or is a sequencing error in the reference sequence.

We found several variants to be unique in line 5B when compared to the remaining five evolved lines after 500 generations of evolution had occurred. Therefore, we were able to back track through frozen stocks to test generations 100, 200, 300, 400, and 500 of line 5B and determined that the transition from sensitivity to resistance of the inhibitory agent occurs between generations 300 and 400 (Figure 2). Sequencing of the populations at generations 300 and 400 then allowed us to narrow the scope of candidate SNPs occurring between these time points.

**Figure 2:**
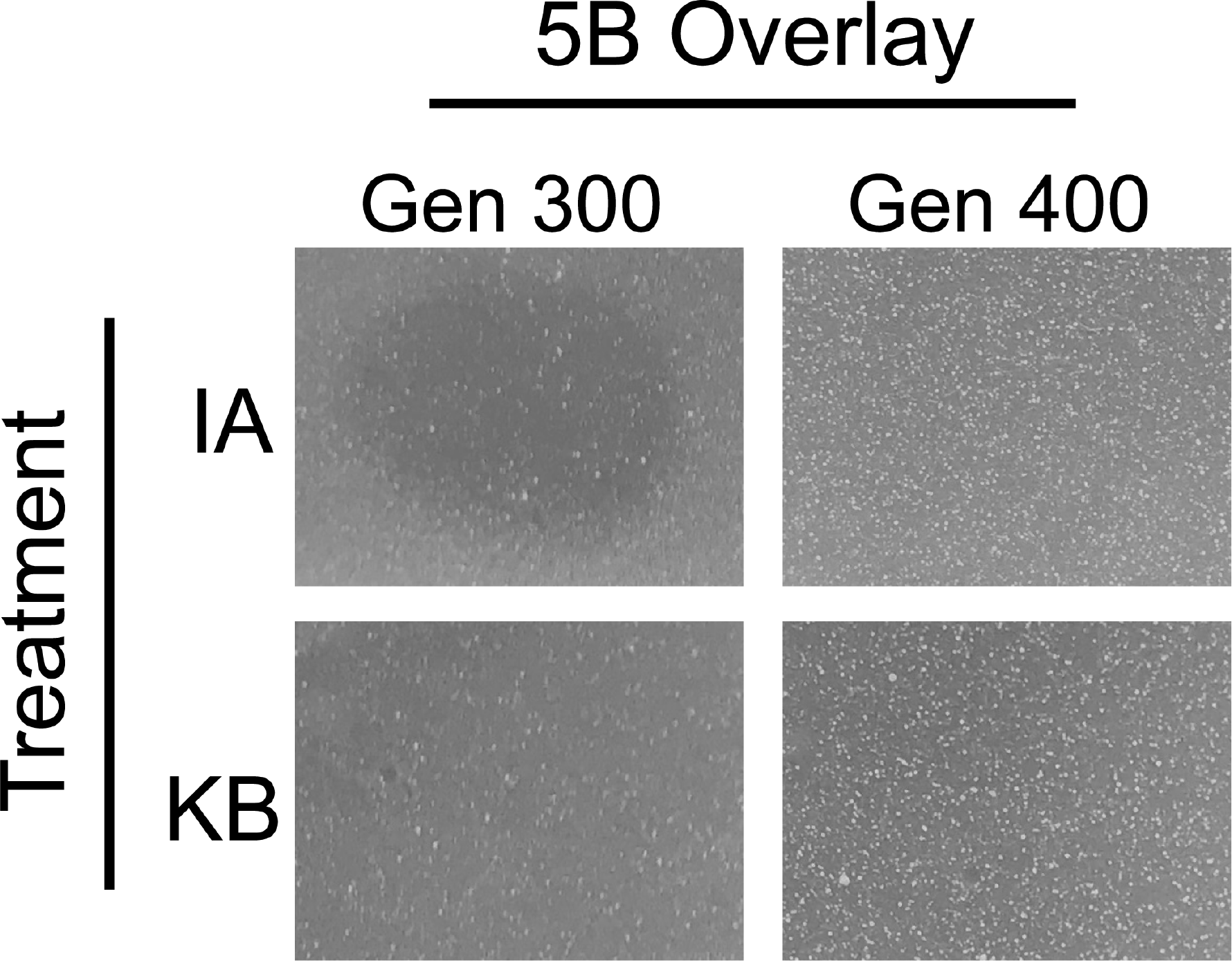
The switch from sensitivity to resistance occurs between generations 300 and 400 in line 5B. We tested generations frozen at various time points of the evolution experiment and found that between generations 300 and 400 line 5B regains resistance to the inhibitory agent produced by *Pseudomonas spp.* A zone of inhibition can be seen when treated with IA (inhibitory agent) at generation 300, and this zone is no longer present at generation 400. Likewise, the negative control (KB media) demonstrates no zones of inhibition as expected. All overlays were plated after 4 hours of growth n KB and spotted with 10μL of *P. stutzeri* filter sterilized supernatants. All images are represented of three biological replicates.

### Conjugation of pMPPla107 from Lines 4B and 5B Results in Resistance to the Inhibitory Agent

The large deletion found in the 4B megaplasmid could alter how the plasmid interacts with its host in a variety of ways including the inhibitory phenotype. Therefore, we hypothesized conjugation of the evolved megaplasmids would transfer resistance to an ancestral strain. Conjugation of evolved pMPPla107 from lines 4B and 5B into a *P. stutzeri* strain containing the ancestral chromosome resulted in resistance to the inhibitory agent while conjugation of ancestral pMPPla107 resulted in sensitivity (Figure 3). Furthermore conjugation of the 5B evolved megaplasmid into *P. syringae* also caused resistance to the inhibitory agent (Supplemental Figure 1). Together these data not only suggest mutations found on pMPPla107 can transfer resistance of the inhibitory agent between *Pseudomonas spp*., but that the underlying mechanism for sensitivity and resistance is shared by Pseudomonads.

**Figure 3:**
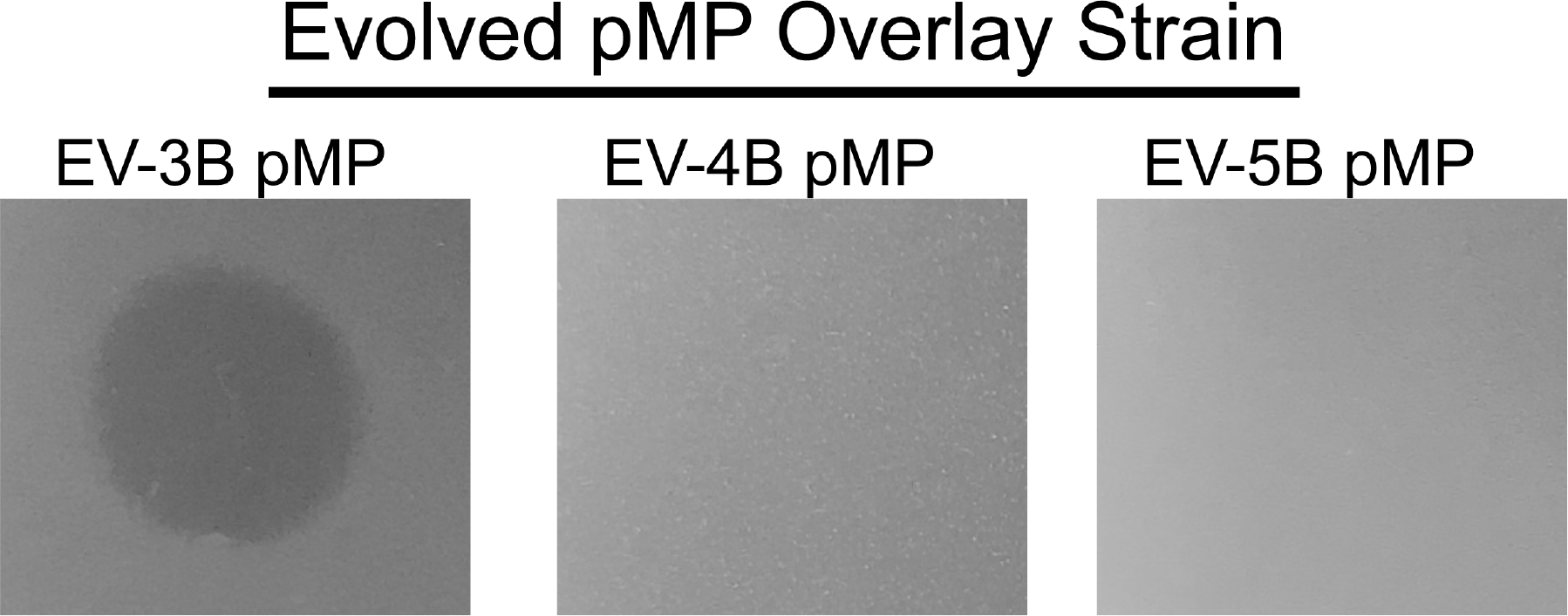
Conjugating evolved pMPPla107 from Lines 4 and 5 into ancestral chromosomal background transfers resistance to the inhibitory agent. Given that Lines 4 and 5 were known to have resistance against the inhibitory agent and chromosomal SNPs did not appear to have any effect we conjugated evolved pMPPla107 into the ancestral chromosomal background (*P. stutzeri* DBL386). We conjugated and used the 3B megaplasmid as a sensitive (positive) control as we knew this evolved strain was still sensitive. When evolved megaplasmids were conjugated into the ancestral background 3B demonstrated sensitivity to the inhibitory agent as expected while 4B and 5B showed resistance. EV = evolved for 500 generations, pMP = pMPPla107. All overlays were plated after 4 hours of growth n KB and spotted with 10μL of *P. stutzeri* filter sterilized supernatants. All images are represented of three biological replicates.

### A 368kb Deletion Occurs in pMPPla107 Evolved Line 4B

Analysis of the genome from isolate DBL408-4BGen500 had the lowest number of variants (20) occurring on the chromosome, while a large deletion of approximately 368kb occurred within pMPPla107 between 131-499kb (Figure 4). This deletion region includes 440 predicted genes without any known homologue and 27 genes with predicted functions (Table 3). Interestingly, there are no repetitive or overlapping sites at the ends of the deletion site suggesting it was not a single deletion event that occurred (https://genomevolution.org/r/uboj). Analysis of previous generations that gave rise to this line against the inhibitory agent indicated that this mutation occurred within the first colony selected for 4B (generation zero) indicating rapid evolutionary changes to pMPPla107 (Figure 5). Although these results indicate that the deletion in line 4B is responsible for resistance to the inhibitory agent, the large size of the deletion and the density of genes within this region make it difficult to discern which gene(s) are responsible for the resistance phenotype in this region.

**Table 3:**
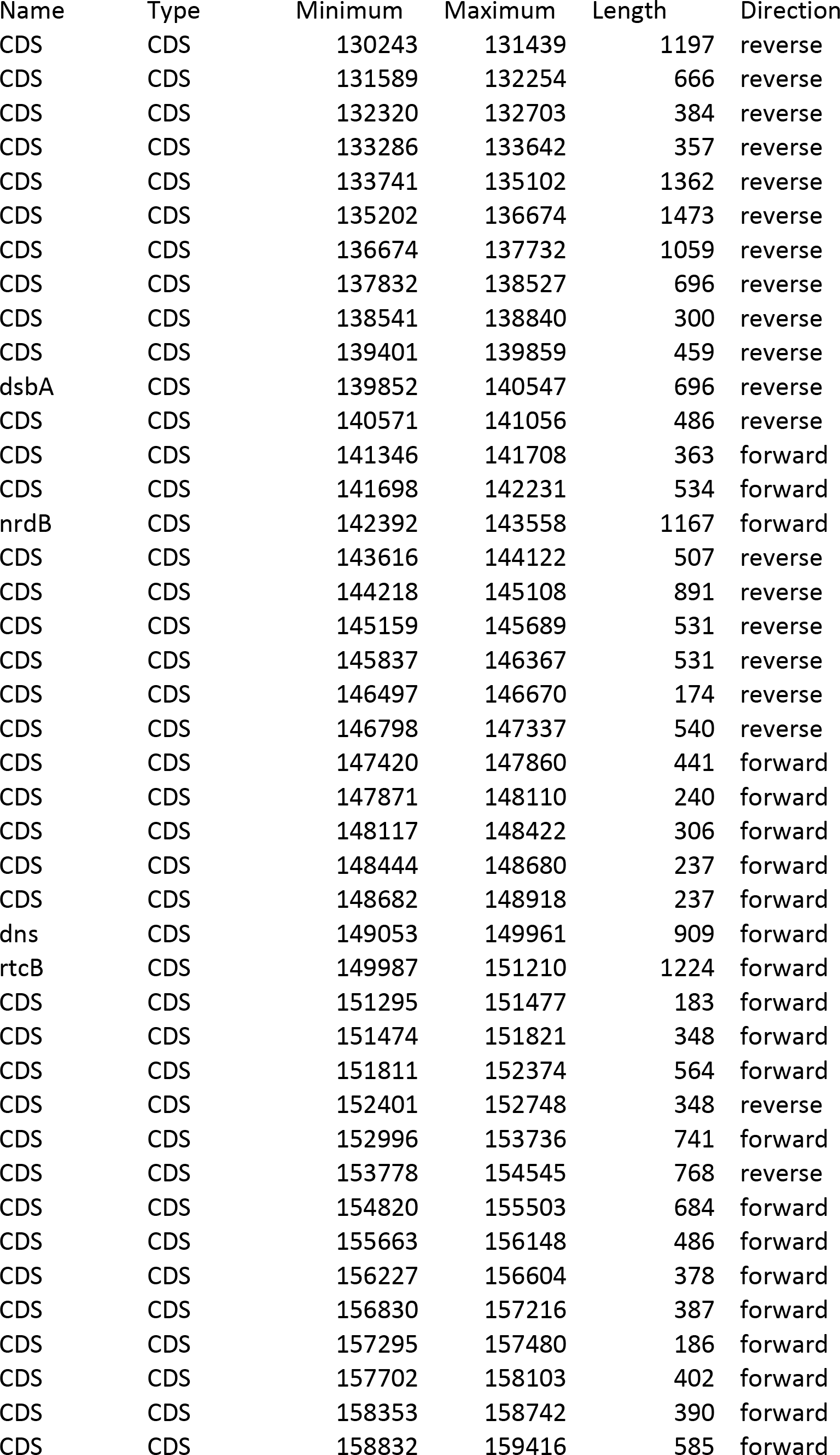
Predicted genes present in the large deletion of pMPPla107 in line 4B.

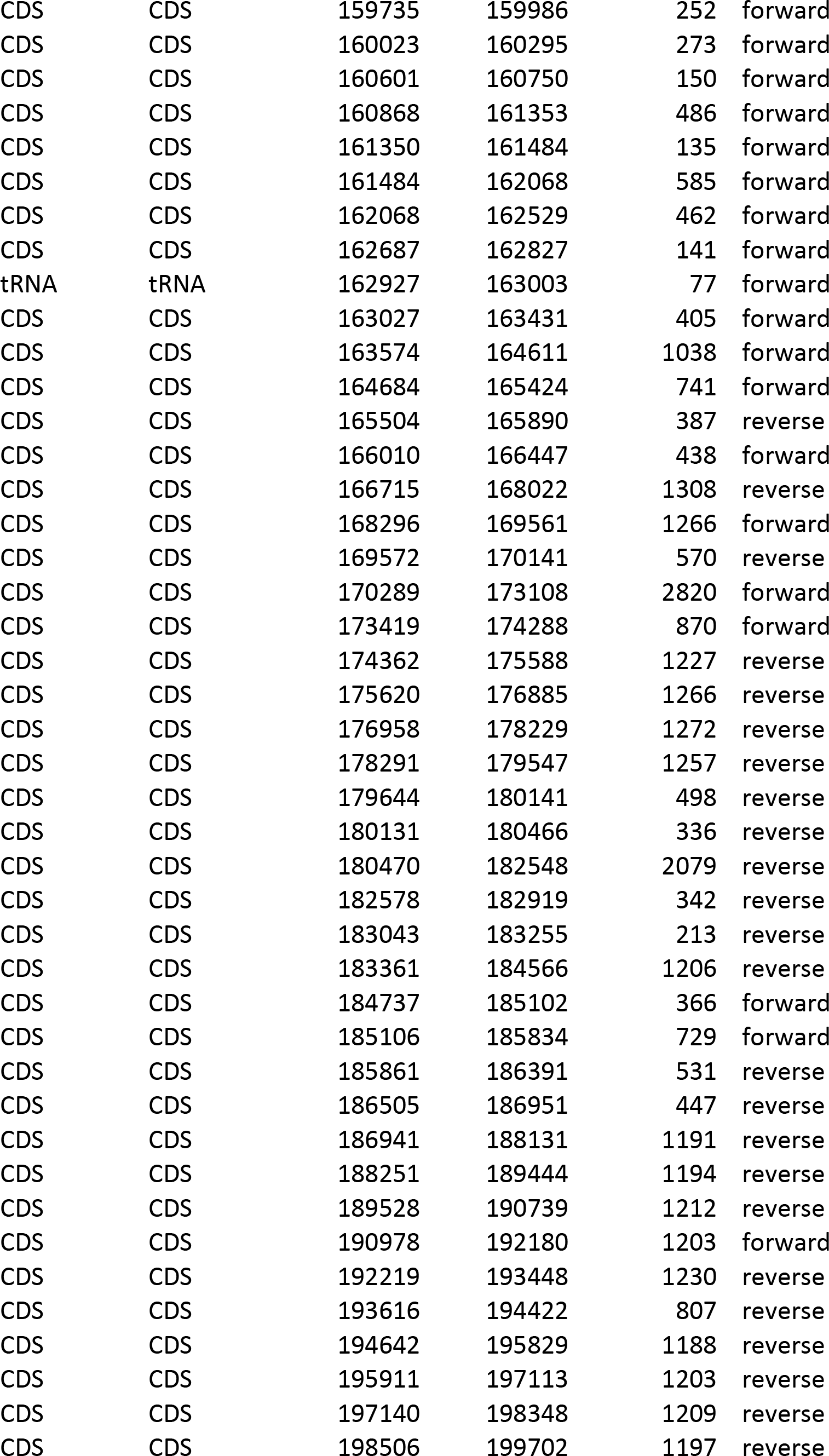

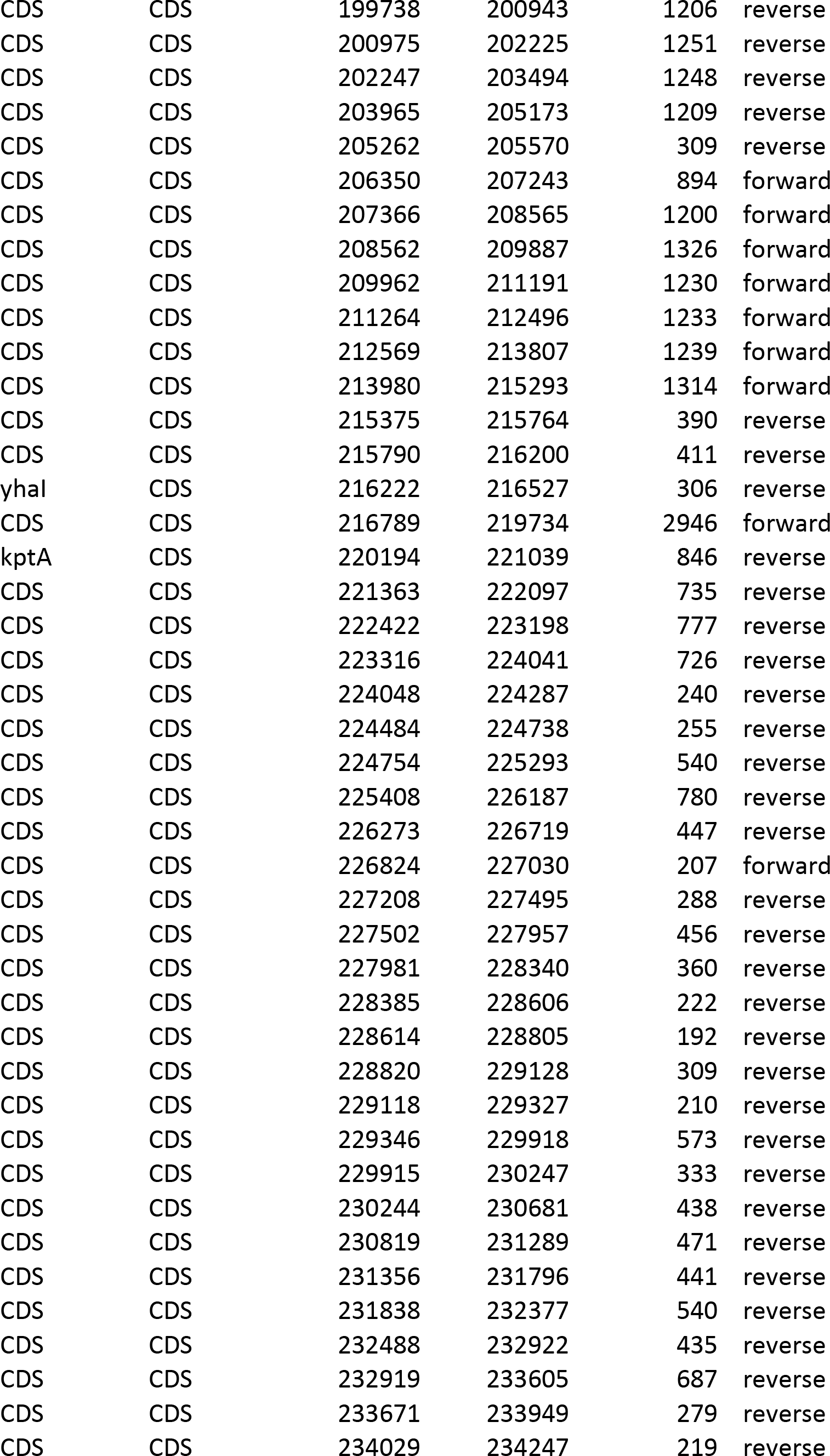

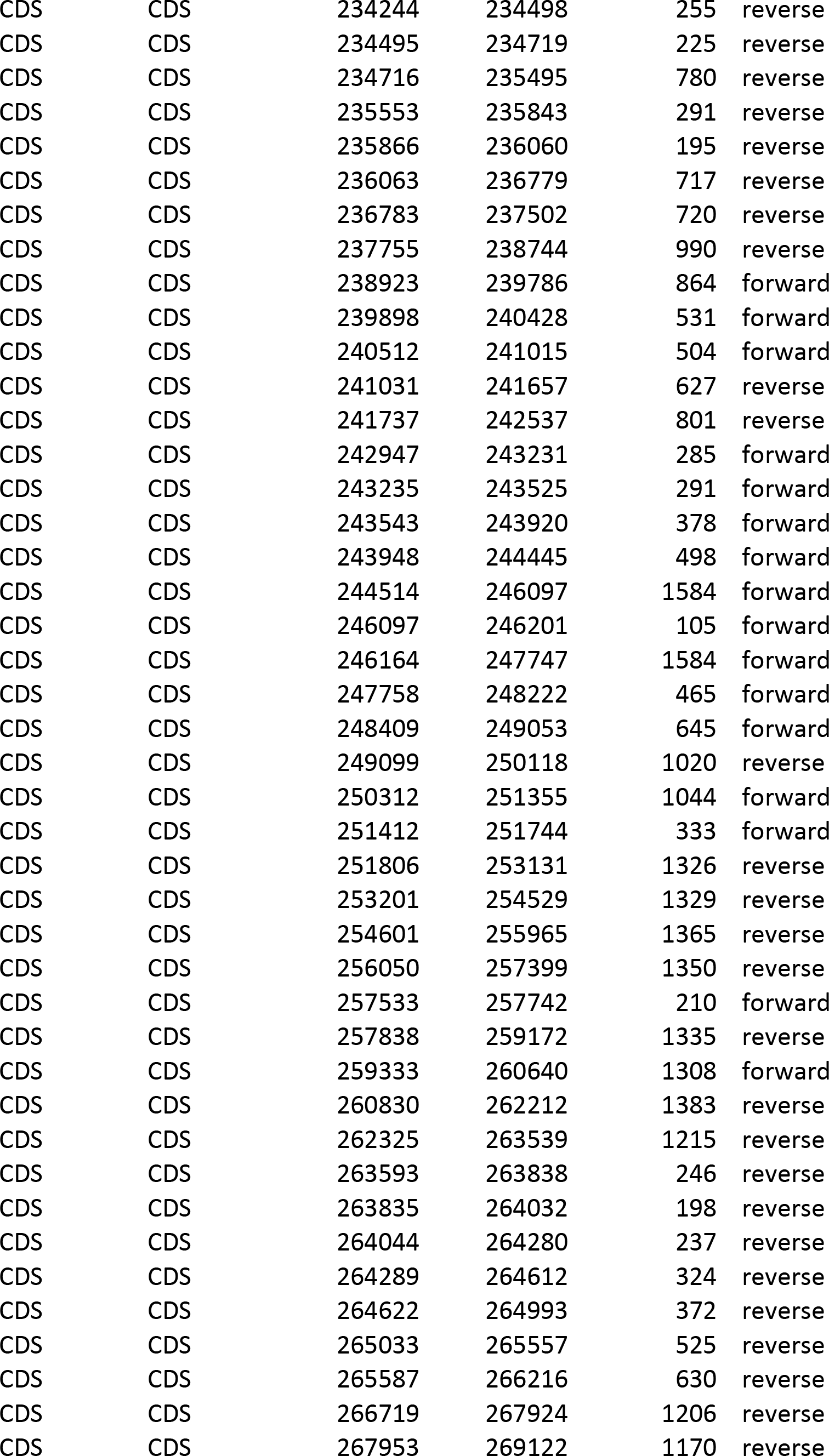

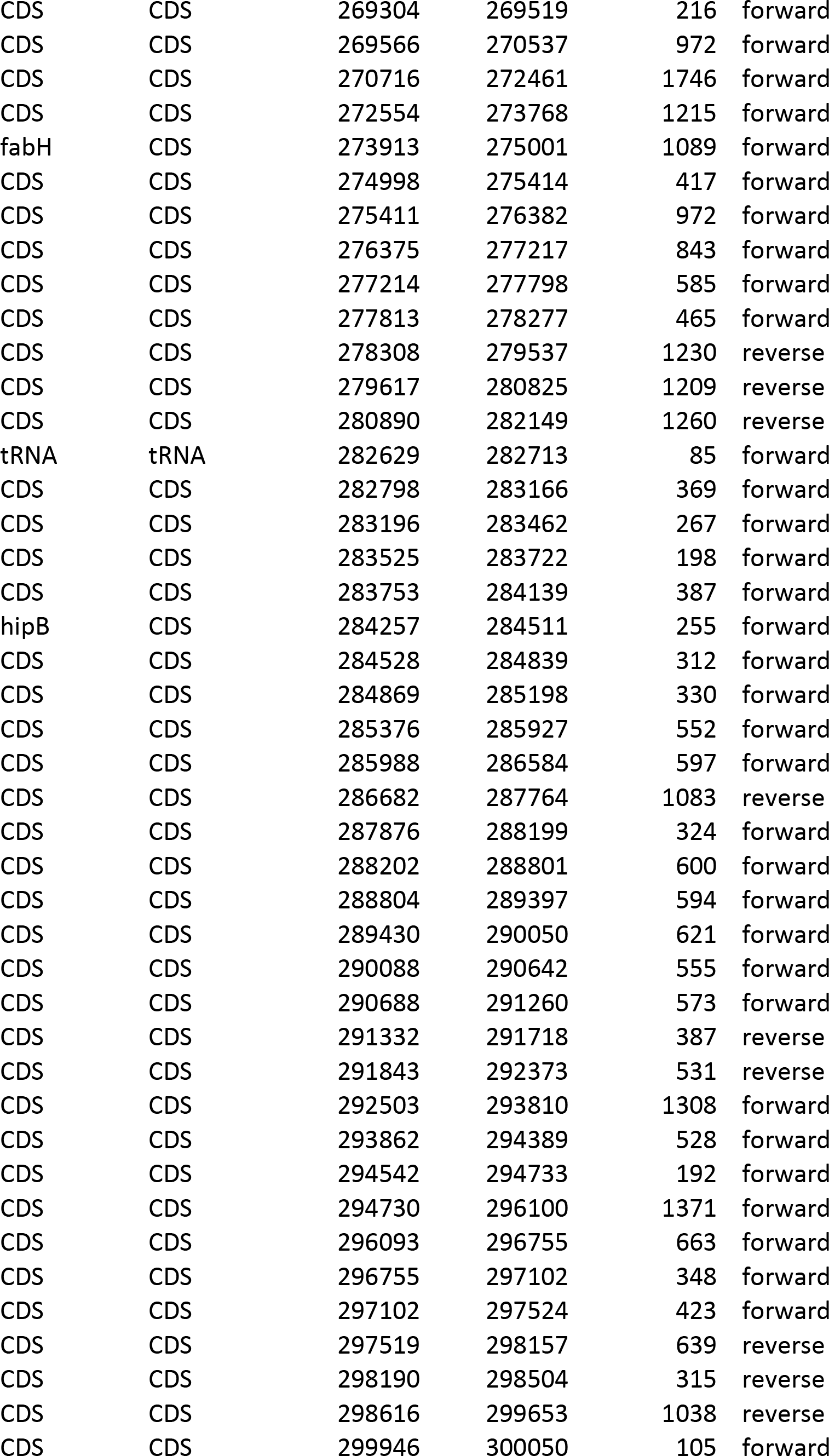

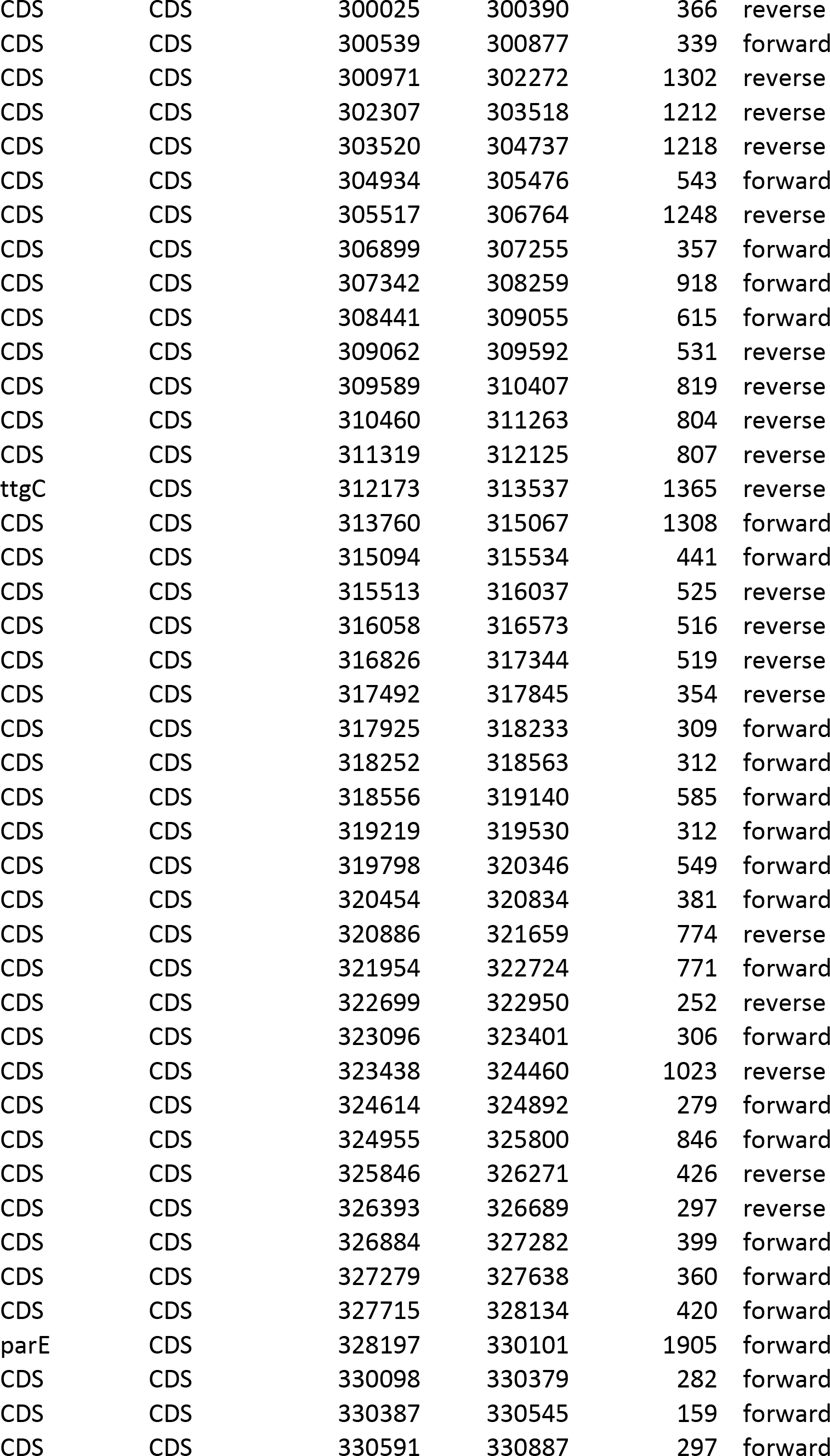

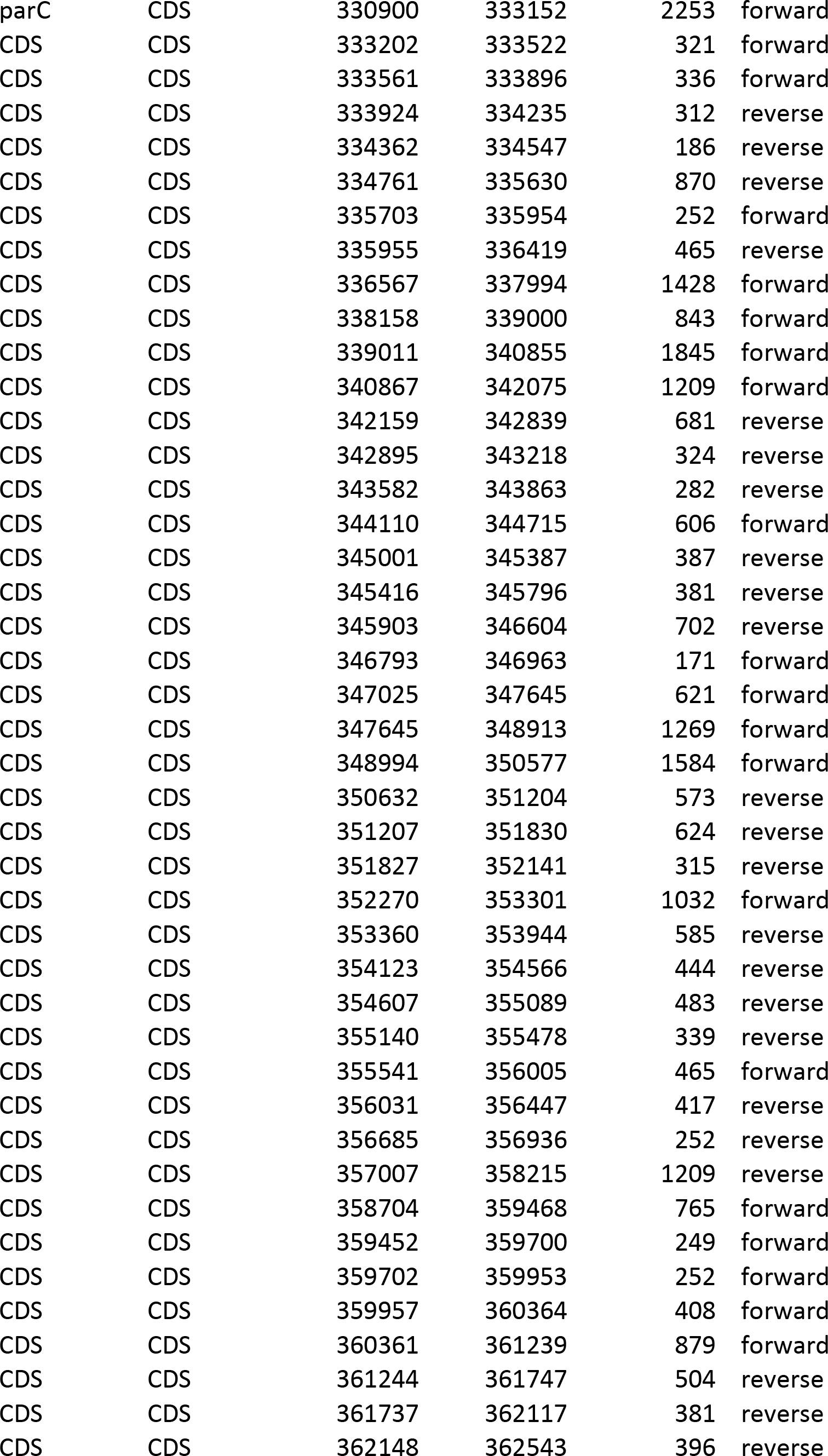

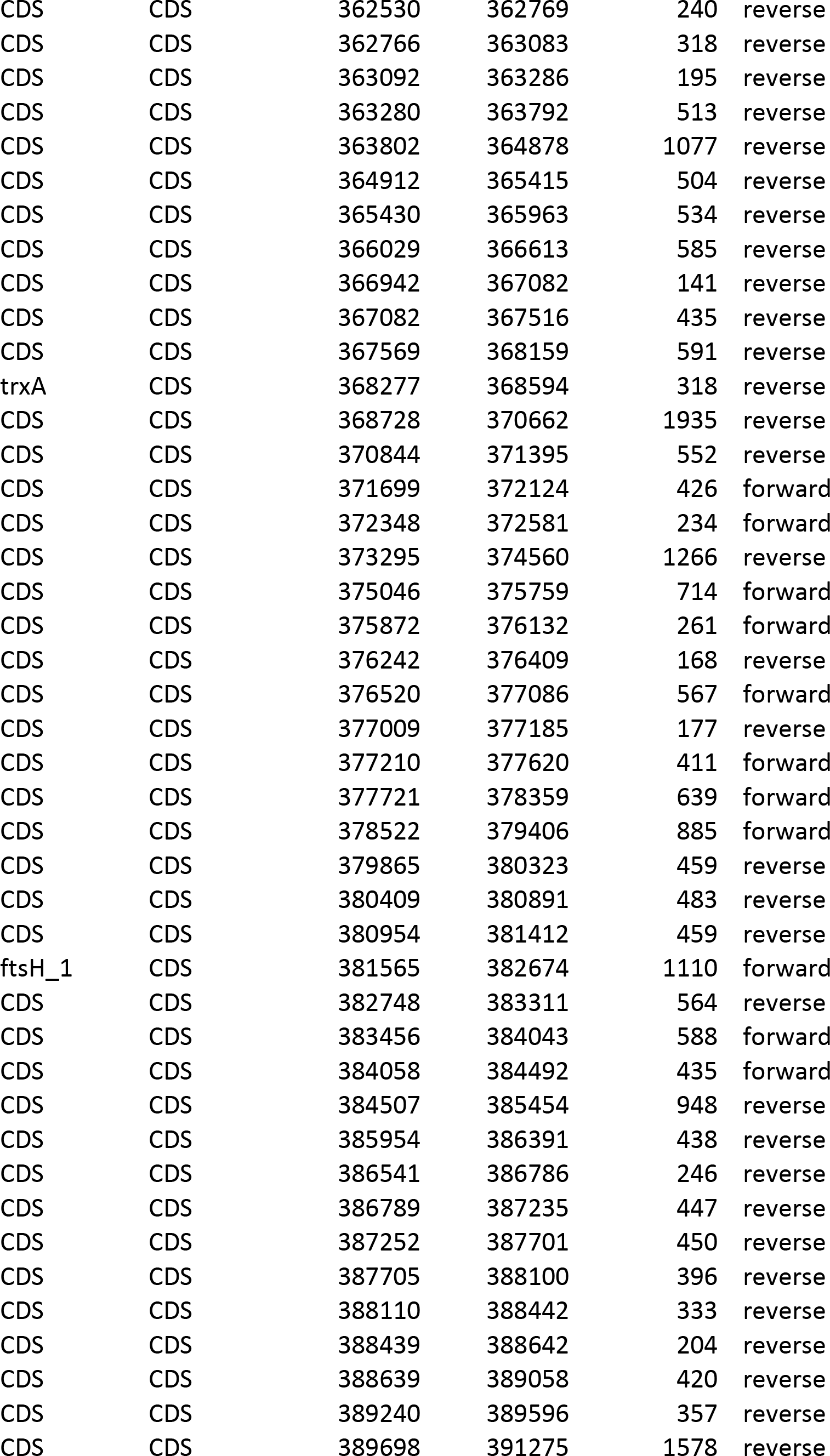

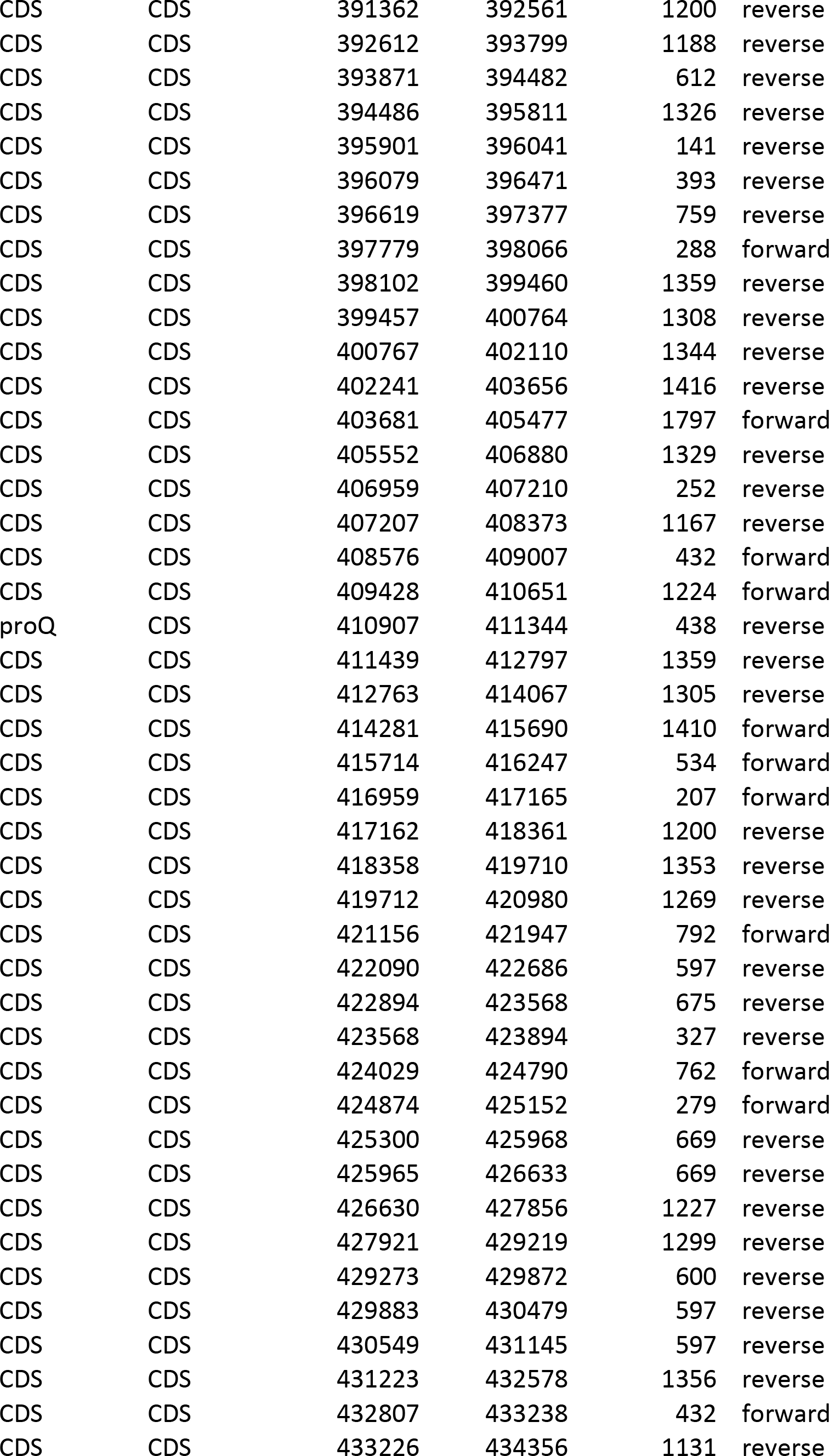

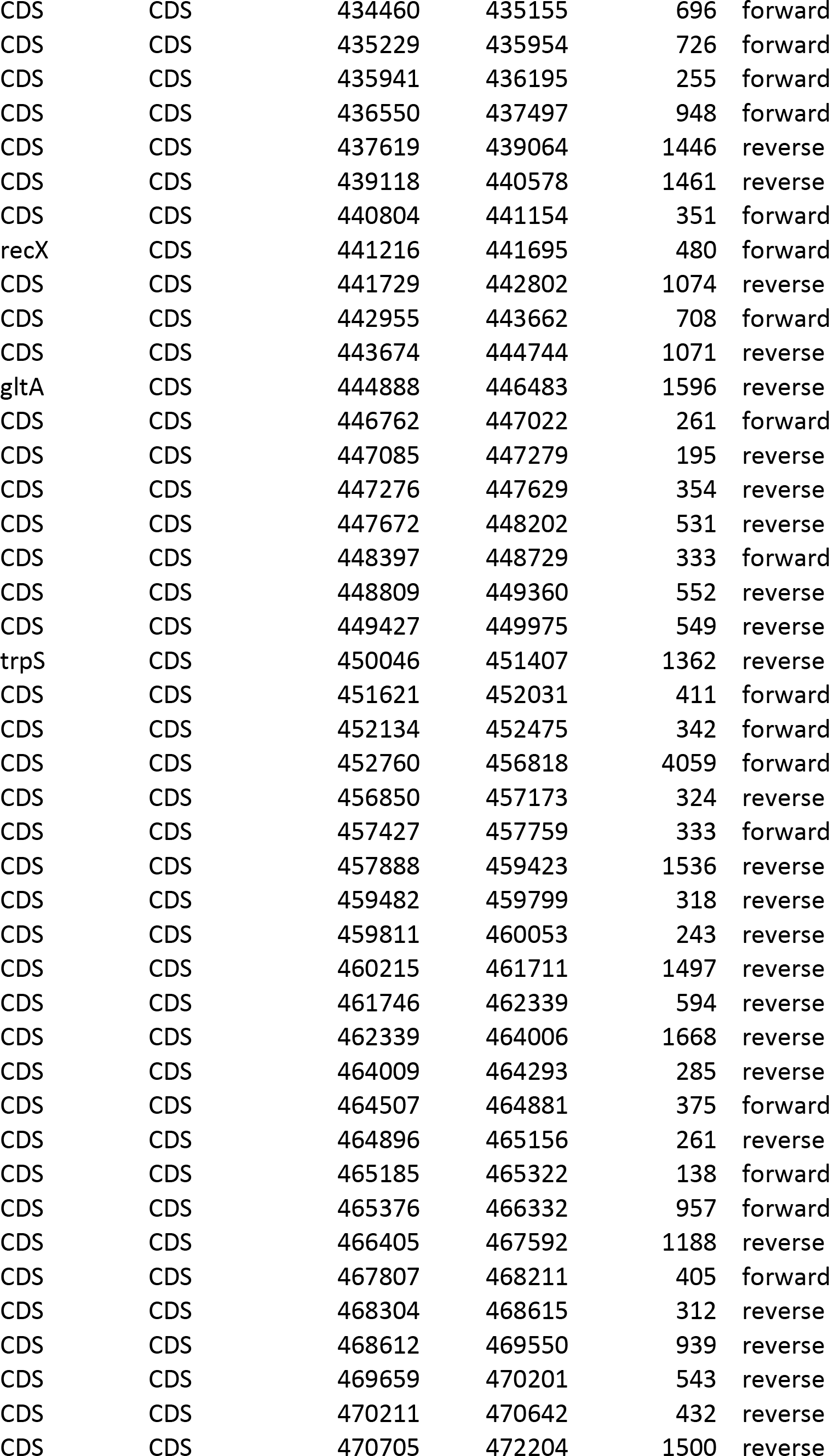

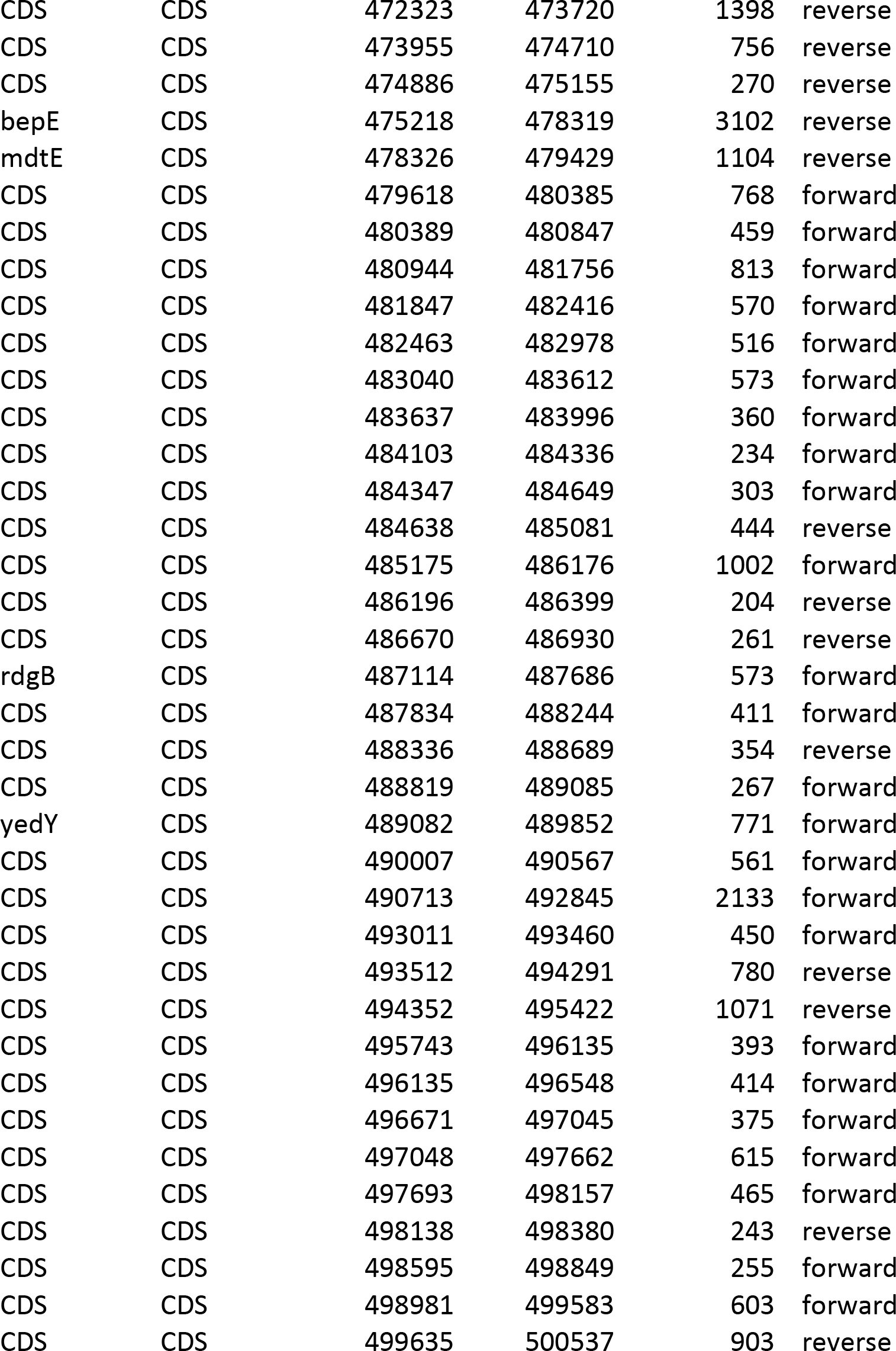

**Figure 4:**
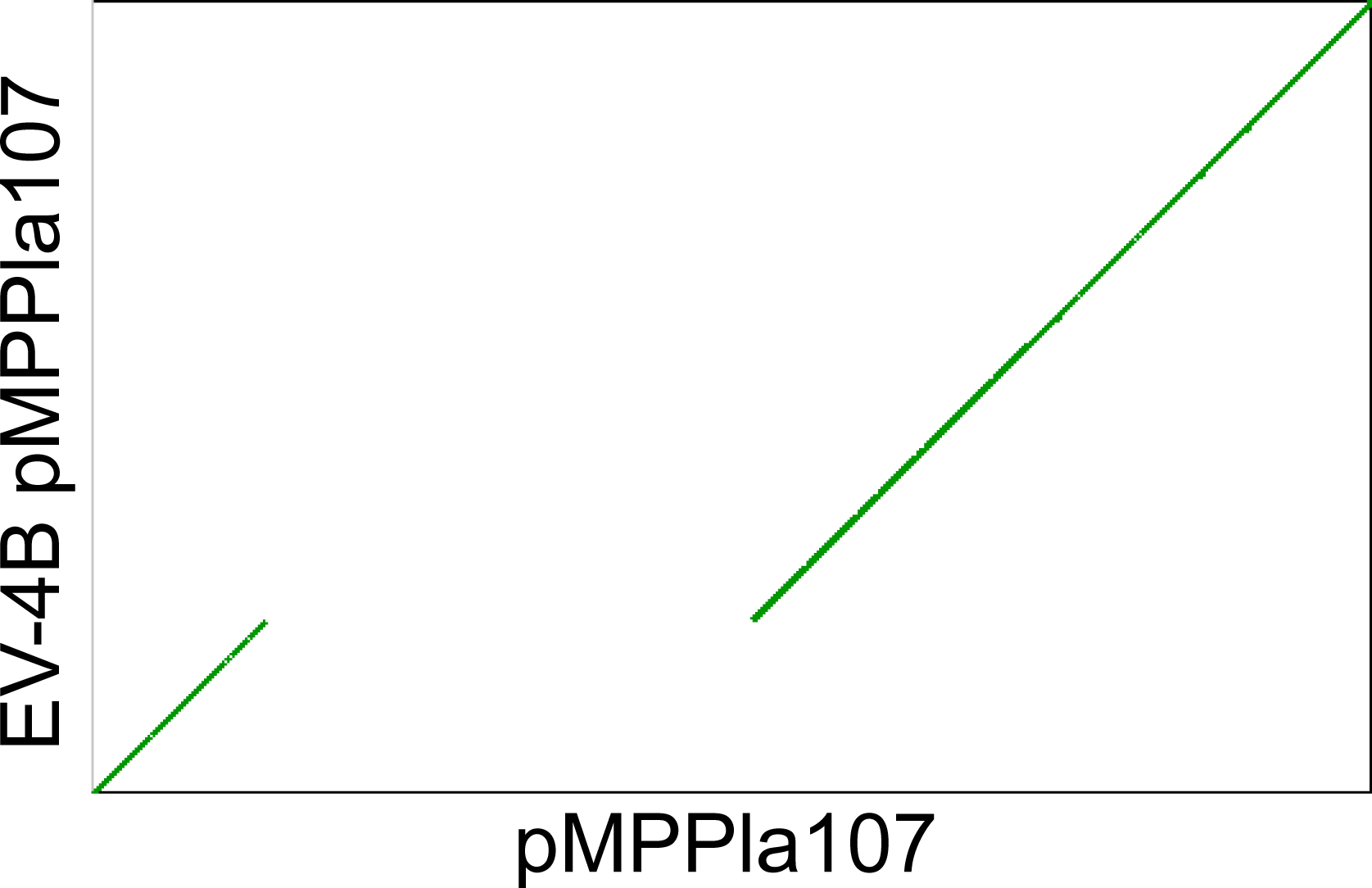
A 368kb occurs in the evolved line 4B megaplasmid. SynMap dotplot visualizes the large deletion occurring from 131-499kb in the evolved 4B pMPPla107 as a large shift across the *x*-axis. The remaining portions of the sequences maintain perfect synteny indicating a clean deletion occurred. The *x*-axis is ancestral pMPPla107 gene order where *x*_1…N_ = gene_1…N_ and the *y*-axis is the line 4B evolved pMPPla107 gene order where *y*_1…N_ = gene_1…N_.

**Figure 5:**
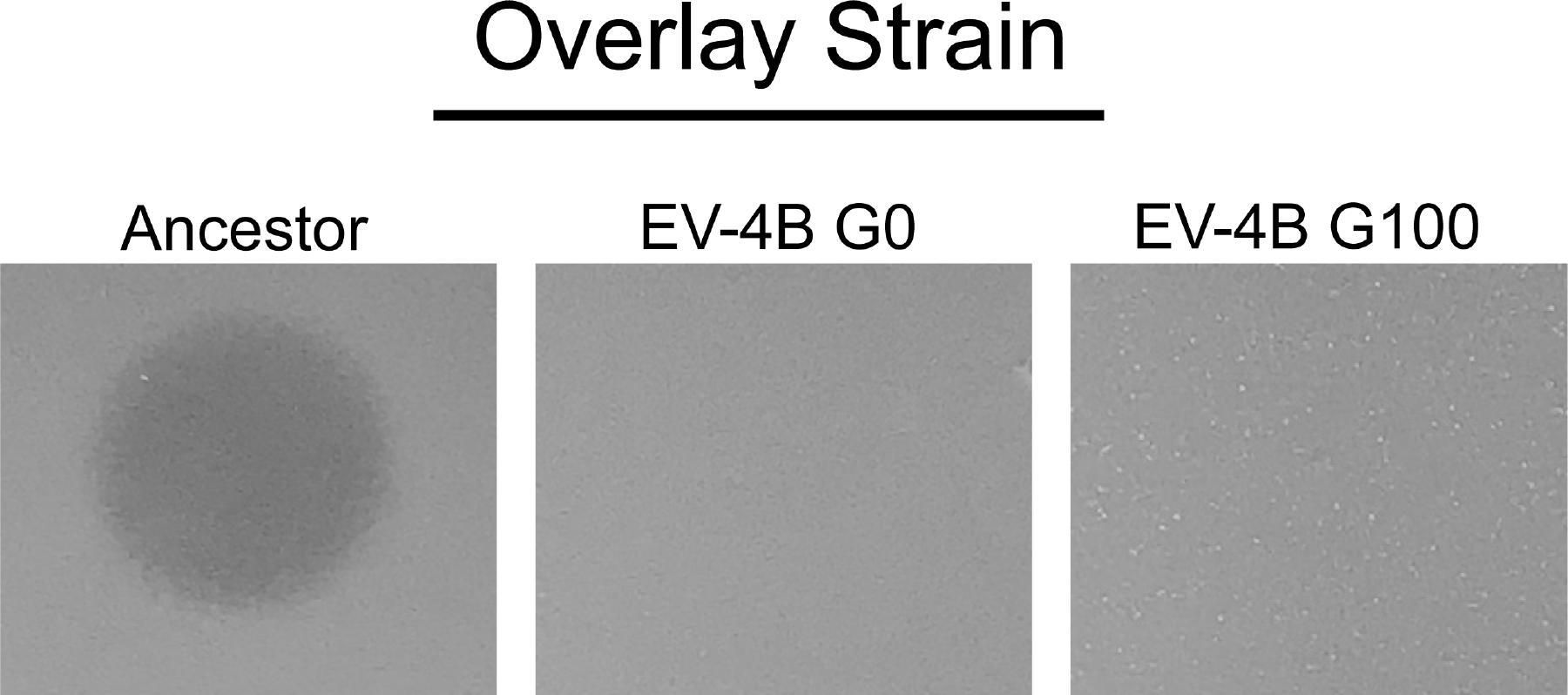
The large deletion in line 4B occurred within the first passage of *P. stutzeri* with pMPPla107. To determine when the deletion occurred we tested frozen generations for sensitivity to the inhibitory agent and found that the deletion was present in the first passage of the evolved line at generation zero. This deletion is maintained in generation 100 (shown) through generation 500 and is the only unique mutation in pMPPla107 other than a synonymous SNP (See Table 2). All overlays were plated after 4 hours of growth n KB and spotted with 10μL of *P. stutzeri* filter sterilized supernatants. All images are represented of three biological replicates.

### A SNP in Line 5B pMPPla107 Causes Resistance to the Inhibitory Agent

Our results indicated that conjugation of pMPPla107 from the evolved 5B line does transfer resistance against the inhibitory agent. Therefore, the variant again, occurs on pMPPla107 (Figure 3). Additionally stated above, we were able to back track through generations of frozen 5B isolates and identified that the resistance phenotype switches from sensitive to resistant between generations 300 and 400 (Figure 2). When comparing SNPs from all six megaplasmids the only unique SNP occurring on pMPPla107 from line 5B between generations 300 and 400 is at 57,137bp and causes a non-synonymous mutation changing a glutamate to a lysine (395 E>K) in an uncharacterized protein. Furthermore, Sanger sequencing of line 5B pMPPla107 in its evolved strain and conjugated to the ancestral *P. stutzeri* strain both confirm the presence of the SNP (<u>DOI: doi.org/10.6084/m9.figshare.7268531</u>). These data suggest that this SNP eliminates the sensitivity phenotype seen by strains that have acquired pMPPla107, thus we name this gene *skaA* for Supernatant Killing Activity. Given that the 5B pMPPla107 SNP occurs outside the deletion region found in the 4B megaplasmid, we also confirm that two separate compensatory strategies exist within pMPPla107 that cause resistance to the inhibitory agent.

## DISCUSSION

We used experimental evolution to identify mutations that are associated with compensation to a unique cost associated with acquisition of megaplasmid pMPPla107. Strains of *P. stutzeri* containing pMPPla107 are sensitized to the presence of a currently unidentified inhibitory agent produced by a variety of Pseudomonas strains under normal growth conditions, and isolates from two of six experimental lines evolve resistance to this inhibition after approximately 500 generations of passage. Numerous studies have found that compensatory mutations to plasmid carriage often occur on the chromosome, but we found that both mutations providing resistance (in lines 4B-500 and 5B-400/5B-500) occur on the megaplasmid.(15–17, 19).

Sequencing of line 4B-500 demonstrated that this line contains a 368kb deletion. This deletion occurs within the same genomic loci of a previously described region of high sequence dissimilarity between the two related plasmids pMPPla107 and pBASL58(5). This suggests a potential cargo region where genes may experience higher mutation and recombination rates resulting in genes that are expendable and provide benefits in certain environments rather than necessary genes for maintenance or transmission. Some of the genes found within this region include efflux pumps, antitoxins, and multidrug resistance proteins all of which may cause resistance to the inhibitory agent (Table 3). It is unclear which of the hundreds of genes in this region is responsible for increased sensitivity to the pseudomonas inhibitory agent, but we identify a specific region, responsible for the sensitivity phenotype.

Conjugating the evolved 5B megaplasmid into an ancestral *P. stutzeri* strain and *P. syringae* demonstrated resistance to the inhibitory agent indicating that the SNP present on pMPPla107-5B was the compensatory mutation and can be transferred across *Pseudomonas spp*. It is still unclear how *skaA* interacts with inhibitory agent or how the 395 E>K SNP changes these interactions. Protein structure and amino acid alignments using Phyre2 and blastx with the NCBI database provided results with low confidence when attempting to identify a function for *skaA* (data not shown)(24, 25).

By combining comparative genomics, microbial genetics and evolutionary methodologies we identified two genetic causes for pMPPla107’s ability to sensitize recipients to a commonly produced inhibitory agent(14). Mutations occurring on the megaplasmid of separately evolved lines indicate that acquisition of pMPPla107 may create conflicts in pseudomonas cellular networks causing a once nontoxic molecule to result in toxicity, but that these mutations alleviate damaged networks. We identify a region on pMPPla107 and a SNP in the gene we now call *skaA* that are responsible for resistance to the pseudomonas inhibitory agent. Our data presented here is the framework on which to begin future work identifying the mechanism behind *skaA* and designing directed deletions within the 4B deletion that will be critical to identifying the other component regarding the inhibitory agent sensitivity phenotype associated with acquisition of pMPPla107.

**Table 1: Variants present in the chromosomes of *P. stutzeri* evolved lines 1-6 after 500 generations of evolution**. The table can be found at Figshare (DOI: doi.org/10.6084/m9.figshare.7393415)

**Table 2: Variants present on pMPPla107 in *P stutzeri* evolved lines 1-6 after 500 generations of evolution**. The table can be found on Figshare (DOI: doi.org/10.6084/m9.figshare.7393493)

## Supplemental Figures

**Supplemental Figure 1:**
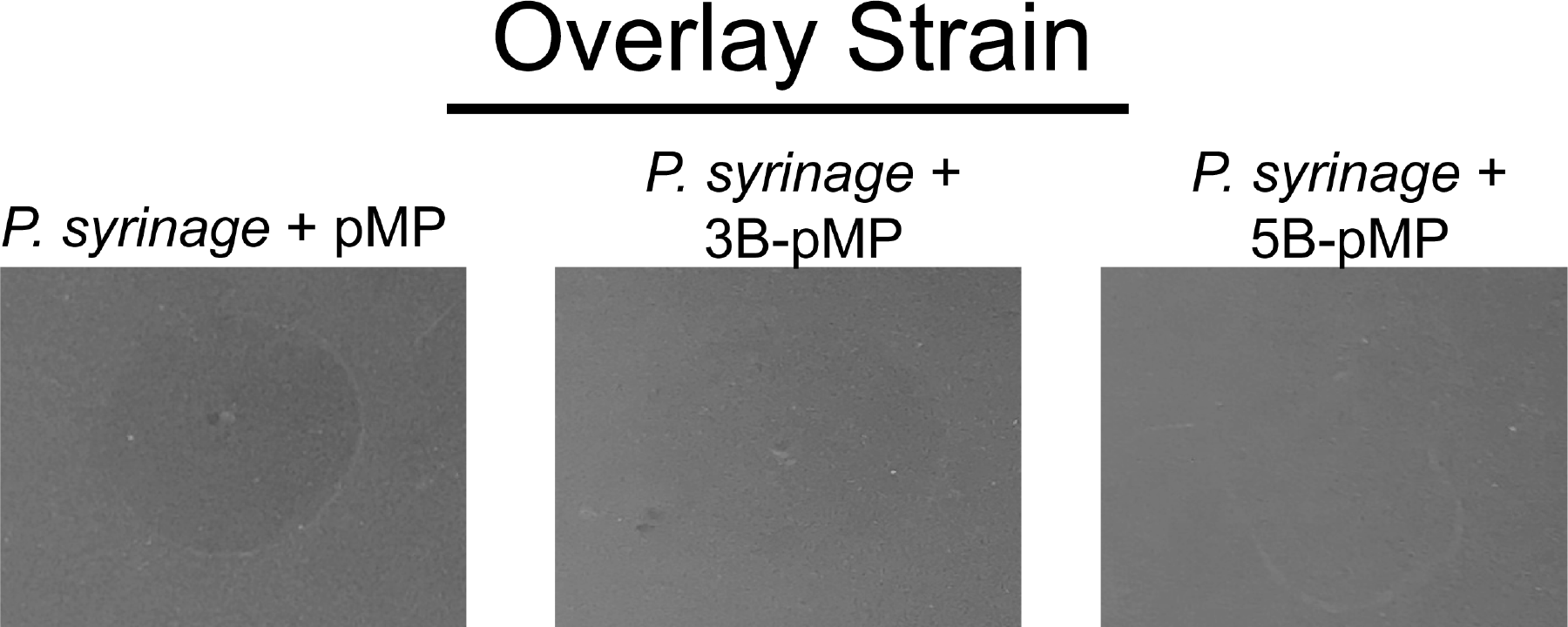
Resistance to the inhibitory agent on the 5B megaplasmid can be transferred to *P. syringae*. Various types of pMPPla107 were conjugated to *P. syringae Pla* YM8003 and then tested for sensitivity to the inhibitory agent on a bacterial overlay. Overlays above are pMP = ancestral pMPPla107, 3B-pMP = evolved line 3B pMPPla107, and 4B-pMP = evolved line 5B pMPPla107. Ancestral and line 3B megaplasmids both transfer sensitivity while line 4B’s megaplasmid transfers resistance to the inhibitory agent. Clearing is less contrasted in overlays with *P. syringae* when compared with *P. stutzeri* overlays due to growth differences (pigment, density) between species. All overlays were plated after 4 hours of growth n KB and spotted with 10μL of *P. stutzeri* filter sterilized supernatants.

## References

1. Kado CI. 1998. Origin and evolution of plasmids. Antonie van Leeuwenhoek 73:117–126.

2. Hülter N, Ilhan J, Wein T, Kadibalban AS, Hammerschmidt K, Dagan T. 2017. An evolutionary perspective on plasmid lifestyle modes. Curr Opin Microbiol 38:74–80.

3. Okubo T, Piromyou P, Tittabutr P, Teaumroong N, Minamisawa K. 2016. Origin and Evolution of Nitrogen Fixation Genes on Symbiosis Islands and Plasmid in Bradyrhizobium. Microbes Environ 31:260–267.

4. Hynes MF, McGregor NF. 1990. Two plasmids other than the nodulation plasmid are necessary for formation of nitrogen-fixing nodules by Rhizobium leguminosarum. Mol Microbiol 4:567–574.

5. Smith BA, Leligdon C, Baltrus D. 2018. Just the Two of Us? A Family of Pseudomonas Megaplasmids Offers a Rare Glimpse Into the Evolution of Large Mobile Elements. bioRxiv 385575.

6. Johnson TJ, Nolan LK. 2009. Pathogenomics of the virulence plasmids of Escherichia coli. Microbiol Mol Biol Rev 73:750–774.

7. San Millan A, MacLean RC. 2017. Fitness Costs of Plasmids: a Limit to Plasmid Transmission. Microbiol Spectr 5.

8. Baltrus DA. 2013. Exploring the costs of horizontal gene transfer. Trends in Ecology & Evolution 28:489–495.

9. Dougherty K, Smith BA, Moore AF, Maitland S, Fanger C, Murillo R, Baltrus DA. 2014. Multiple phenotypic changes associated with large-scale horizontal gene transfer. PLoS ONE 9:e102170.

10. Romanchuk A, Jones CD, Karkare K, Moore A, Smith BA, Jones C, Dougherty K, Baltrus DA. 2014. Bigger is not always better: transmission and fitness burden of ∼1MB Pseudomonas syringae megaplasmid pMPPla107. Plasmid 73:16–25.

11. Glick BR. 1995. Metabolic load and heterologous gene expression. Biotechnol Adv 13:247–261.

12. Sato T, Kuramitsu H. 1998. Plasmid maintenance renders bacteria more susceptible to heat stress. Microbiol Immunol 42:467–469.

13. Heuer H, Fox RE, Top EM. 2007. Frequent conjugative transfer accelerates adaptation of a broad-host-range plasmid to an unfavorable Pseudomonas putida host. FEMS Microbiology Ecology 59:738–748.

14. Smith B, Feinstein Y, Clark M, Baltrus D. 2019. A Moving Target: The Megaplasmid pMPPla107 Sensitizes Cells to an Inhibitory Agent Conserved Across Pseudomonas spp. bioRxiv 537589.

15. Harrison E, Guymer D, Spiers AJ, Paterson S, Brockhurst MA. 2015. Parallel compensatory evolution stabilizes plasmids across the parasitism-mutualism continuum. Curr Biol 25:2034–2039.

16. Loftie Eaton W, Bashford K, Quinn H, Dong K, Millstein J, Hunter S, Thomason MK, Merrikh H, Ponciano JM, Top EM. 2017. Compensatory mutations improve general permissiveness to antibiotic resistance plasmids. Nat Ecol Evol 1:1354–1363.

17. Yano H, Wegrzyn K, Loftie Eaton W, Johnson J, Deckert GE, Rogers LM, Konieczny I, Top EM. 2016. Evolved plasmid-host interactions reduce plasmid interference cost. Mol Microbiol.

18. Bouma JE, Lenski RE. 1988. Evolution of a bacteria/plasmid association. Nature 335:351–352.

19. Morton ER, Merritt PM, Bever JD, Fuqua C. 2013. Large deletions in the pAtC58 megaplasmid of Agrobacterium tumefaciens can confer reduced carriage cost and increased expression of virulence genes. Genome Biol Evol 5:1353–1364.

20. Sikorski J, Teschner N, Wackernagel W. 2002. Highly different levels of natural transformation are associated with genomic subgroups within a local population of Pseudomonas stutzeri from soil. Appl Environ Microbiol 68:865–873.

21. Seemann T. 2014. Prokka: rapid prokaryotic genome annotation. Bioinformatics 30:2068–2069.

22. Haug-Baltzell A, Stephens SA, Davey S, Scheidegger CE, Lyons E. 2017. SynMap2 and SynMap3D: web-based whole-genome synteny browsers. Bioinformaticsedn. 33:2197–2198.

23. Baltrus DA, Nishimura MT, Romanchuk A, Chang JH, Mukhtar MS, Cherkis K, Roach J, Grant SR, Jones CD, Dangl JL. 2011. Dynamic Evolution of Pathogenicity Revealed by Sequencing and Comparative Genomics of 19 Pseudomonas syringae Isolates. PLOS Pathog 7:e1002132.

24. Kelley LA, Mezulis S, Yates CM, Wass MN, Sternberg MJE. 2015. The Phyre2 web portal for protein modeling, prediction and analysis. Nat Protoc 10:845–858.

25. Altschul SF, Gish W, Miller W, Myers EW, Lipman DJ. 1990. Basic local alignment search tool. Journal of Molecular Biology 215:403–410.

26. Carroll AC, Wong A. 2018. Plasmid persistence: costs, benefits, and the plasmid paradox. Can J Microbiol 64:293–304.

27. San Millan A, Peña-Miller R, Toll-Riera M, Halbert ZV, McLean AR, Cooper BS, MacLean RC. 2014. Positive selection and compensatory adaptation interact to stabilize non-transmissible plasmids. Nature Communications 5:5208.

